# Synergistic interactions between detritivores disappear under reduced rainfall

**DOI:** 10.1101/2020.09.29.318592

**Authors:** François-Xavier Joly, Euan McAvoy, Jens-Arne Subke

## Abstract

Understanding the consequences of altered rainfall patterns on litter decomposition is critical to predicting the feedback effect of climate change on atmospheric CO_2_ concentrations. While their effect on microbial decomposition received considerable attention, their effect on litter fragmentation by detritivores, the other dominant decomposition pathway, remains largely unexplored. Particularly, it remains unclear how different detritivore species and their interactions respond to changes in rainfall quantity and frequency. To fill this knowledge gap, we determined the contribution to litter decomposition of two detritivore species (millipede and isopod), separately and in combination, under contrasting rainfall quantity and frequency in a temperate forest. Although halving rainfall quantity and frequency decreased top-soil moisture by 7.8 and 13.1%, respectively, neither millipede- nor isopod-driven decomposition were affected by these changes. In contrast, decomposition driven by both detritivore species in combination was 65.5% higher than expected based on monospecific treatments under high rainfall quantity, but unchanged or even lower under low rainfall quantity. This indicates that while detritivore activity is relatively insensitive to changes in rainfall patterns, large synergistic interactions between detritivore species may disappear under future rainfall patterns. Incorporating interspecific interactions between decomposers thus seems critical to evaluate the sensitivity of decomposition to altered rainfall patterns.

## Introduction

Understanding the consequences of climate change for processes underpinning the global carbon cycle is essential to predicting its feedback effect on atmospheric CO_2_ concentrations. This critical challenge prompted major research efforts evaluating the response of plant litter decomposition – the largest flux of CO_2_ to the atmosphere (Schlesinger 2005) – to expected changes in climate change. In particular, predictions of altered rainfall patterns with larger but less frequent rainfall events leading to enhanced probabilities of drought and heavy rains (IPCC 2013) led numerous studies to evaluate how decomposition responds to various rainfall manipulation. To date, the vast majority of these studies have focused on the response of microbial decomposition (Sanaullah et al. 2012, Walter et al. 2013, Joly et al. 2017), and reported strong negative responses of microbial decomposition to reduced rainfall quantity and frequency. In contrast, the response of detritivorous soil animals to altered rainfall patterns remains poorly documented.

Litter-feeding soil invertebrates (detritivores hereafter) play a major role in organic matter turnover by consuming large amounts of decomposing litter (García-Palacios et al. 2013) and returning most of it to the soil as feces (David 2014). This conversion of litter into feces consisting of a myriad of fecal particles facilitates further leaching and microbial activity, thereby accelerating decomposition (Joly et al. 2020). Despite the importance of this pathway, its response to expected changes in rainfall patterns is difficult to predict. Particularly, it remains poorly understood how detritivores respond to changes in rainfall quantity (cumulative rainfall) and frequency (infrequent large rainfall event vs frequent smaller events). Among the few studies that evaluated this response, Coulis et al. (2013) reported a limited decrease in the feeding activity of one millipede species (*Ommatoiulus sabulosus*) in response to reduced rainfall quantity and frequency. In turn, Joly et al. (2019) found that the feeding activity of one isopod species (*Armadillidium vulgare*) was not affected by large changes in rainfall quantity, but controlled by rainfall frequency with higher activity at low rainfall frequency. The relative insensitivity of detritivore activity to reduced rainfall quantity suggests a desiccation resistance, either morphological (e.g., exoskeleton; low surface/volume ratio) or behavioral (e.g., mobility allowing to shelter or find water in moist soil areas). In turn, the higher activity at lower rainfall frequency reported by Joly et al. (2019) suggests that, for a given rainfall quantity, large and infrequent rainfall events trigger detritivore activity more efficiently than more frequent but smaller events, as hypothesized by Nielsen and Ball (2015), or that alternation between dry and moist conditions induces compensatory feeding, with detritivores consuming higher quantities of moist litter following drought periods to satisfy their water requirements. The contrasting responses to rainfall frequency between the two studies indicate that the influence of rainfall pattern on detritivore-driven decomposition may depend on detritivore species, yet species-specific sensitivities to rainfall patterns are unclear. Also, both studies were performed under laboratory conditions with detritivores collected from populations commonly exposed to drought (Mediterranean shrubland and dryland, respectively) and there are so far no data on the importance of this rainfall pattern control under field conditions and in mesic ecosystems.

In addition to interspecific differences in the response of feeding activity to rainfall pattern, the contribution of detritivores to decomposition also depends on the nature of interspecific interactions between cooccurring detritivores (Gessner et al. 2010). Several studies reported that when different soil animal species co-occur, their contribution to decomposition can be larger than predicted based on monospecific treatments (Heemsbergen et al. 2004, Hedde et al. 2010, DeOliveira et al. 2010). These synergistic effects of detritivore diversity may contribute substantially to decomposition, but the response of these interactions to climatic change is largely unknown. Theory predicts that as environmental stress increases, interspecific interactions switch from competition to complementarity and facilitation (Bertness and Callaway 1994). This stress-gradient hypothesis implies that an increase in environmental stress could drive a switch from antagonistic or non-additive diversity effects on ecosystem functioning to synergistic effects. The hypothesis is relatively well supported for plant-plant interactions and associated primary productivity (He et al. 2013), but remains poorly considered for interactions between soil organisms and associated decomposition processes, with the rare studies focusing on interactions between detritivore diversity and rainfall pattern reporting contrasted results (Collison et al. 2013, Coulis et al. 2015). Examining the nature of interactions between detritivores and their response to altered rainfall patterns thus appears critical to predict decomposition under future climate scenarios.

We evaluated the consequences of halving rainfall quantity and frequency on the feeding activity of two common detritivore species from distant phylogenetic groups (millipedes and isopods), and on their interactions. We hypothesized that (H1) both detritivore species are more strongly affected by reductions in rainfall frequency than quantity, and that (H2) synergistic interactions between detritivore species increase with reduced rainfall quantity and frequency. We tested these hypotheses by measuring the contribution of millipedes (*Glomeris marginata*) and isopods (*Armadillidium vulgare*), separately and in combination, to leaf litter decomposition during a six-week field experiment in a Scottish temperate forest during which we fully manipulated rainfall patterns.

## Methods

We conducted this experiment in a mixed woodland of the University of Stirling (56°08’34” N, 3°55’12” W; 40 m a.s.l), located within the Scottish Lowlands, UK. The climate is characterized by mean annual precipitation of 1019 mm and a mean annual temperature of 8.5 °C (MetOffice, 1981-2010). The months of August and September, during which we ran this experiment, have mean monthly precipitation of 72.4 and 90.6 mm, respectively and have 11.4 and 11.9 days with rainfall > 1 mm, respectively. The vegetation at the site consists in a mixture of broadleaf and coniferous tree species, including *Acer pseudoplatanus* L., *Fagus sylvatica* L., *Aesculus hippocastanum* L., *Picea sitchensis* (Bong.) Carr, and *Larix decidua* Mill. In May 2018, we collected two detritivore species from nearby woodlands, including the millipede *Glomeris marginata* (Villers, 1789) in a woodland near Peebles, Scotland, UK (55°38’46” N, 3°07’55” W), and the isopod *Armadillidium vulgare* (Latreille, 1804) in a coastal woodland near Dunfermline, Scotland, UK (56°01’35”N, 3°23’14”W). We chose these species as they are widespread across Europe, occurring in diverse ecosystems from Mediterranean to temperate. Detritivores were kept in plastic boxes and fed with moist litter until use. In July 2018, we collected *Aesculus hippocastanum* leaf litter from the study site. This litter consisted in 0.465 g C g^-1^ litter and 0.0089 g N g^-1^ litter, for a C:N ratio of 52.0. Litter of this tree species is characterized as relatively recalcitrant, with high C:N ratio and tannin concentrations, and low concentration of water-soluble compounds and capacity to retain water (Joly et al. 2020). We used decomposing leaf litter rather than freshly senesced litter because detritivores prefer feeding on partially decomposed litter (David and Gillon, 2002). We air-dried the litter, cleaned it of debris (twigs, non-targeted litter species), and stored it in paper bags until the start of the experiment.

We manipulated rainfall quantity (18 vs. 36 mm/month), rainfall frequency (one vs. two pulses a week) and detritivore community composition (control without detritivore; millipedes only; isopods only; millipedes and isopods together) in a full-factorial design of field mesocosms during a six-week period in August and September 2018. Each treatment combination was replicated five times for a total of 80 mesocosms (2 rainfall quantities × 2 rainfall frequencies × 4 detritivore treatments × 5 replicates). To do so, we set up five replicate full-rainout shelters (6.25 m^2^ surface area), consisting of 2.5 × 2.5 m wooden frames covered by a transparent polythene sheet (0.127 mm thick, 508 g/m^2^), elevated at 0.75 m above the soil surface with wooden legs in corners to allow air passage. To prevent rainwater from accumulating in the middle of the polythene sheet, we placed a 1 m stake in the center of each shelter to elevate the sheet and allow the rainwater to runoff. Under each shelter, we set up four subplots (0.5 m^2^ surface area); one per rainfall treatment. These subplots consisted in 0.7 × 0.7 m wooden frames placed on the soil surface and inserted 5 cm into the soil. We placed these frames such that they were 40 cm away from the shelter edges to avoid moisture from shelter run-off, and 20 cm from one another. Within subplots, we removed the existing litter layer and inserted four mesocosms (one per detritivore treatment) made of PVC pipe (9 cm long; 16 cm diameter) 7 cm into the soil. Because detritivores shelter during dry conditions to avoid desiccation, we added small tunnels made of PVC pipe (35 mm long; 35 mm diameter) buried halfway, lengthwise, in the middle of each mesocosm. We then added 6.0 ± 0.01 g of air-dried leaf litter to each mesocosm. We converted air-dried litter mass into 60°C dry mass by weighing air-dried litter subsamples, drying them at 60 °C for 48 h, and reweighing them to obtain dry mass. In each subplot, we randomly attributed a detritivore treatment to mesocosms and added either (1) no detritivore for the control treatment without detritivore, or 0.6 ± 0.01 g of (2) millipedes, (3) woodlice, or (4) millipedes and woodlice. Because of variation in body size, the number of detritivores per microcosm varied between 5 and 6. To prevent detritivores from escaping while allowing evaporation, we covered mesocosms with a 2 × 2 mm nylon mesh secured with rubber bands. Within each subplot, we also added leaf litter between mesocosms, in a similar density as within mesocosms to ensure homogeneous surface conditions (including desiccation) throughout the subplot. We randomly attributed a rainfall treatment to each subplot for each rainout shelter. Rainfall quantity and frequency treatments consisted in 18 and 36 mm/month, with the equivalent weekly amounts added as large pulses once a week or smaller pulses twice a week. These quantities approximately reflected the actual throughfall (ca. 42 mm/month) during the incubation period and a 50% reduction (ca. 21 mm/month), respectively. This throughfall was estimated by correcting the mean monthly rainfall by an estimated 50% rainfall losses due to canopy interception and stemflow (Johnson 1990). Similarly, the frequencies approximately reflected the actual number of days with > 1 mm of throughfall and a 50% reduction, respectively. We added the water pulses for both frequency treatments on Monday mornings and those for the high frequency treatment additionally on Thursday afternoons. These pulses were applied to mesocosms by watering subplots with a Knapsack pressure sprayer (PS16, Kingfisher). We monitored the moisture of the topsoil (top 1 cm) throughout the incubation by collecting soil samples from all subplots on Mondays and Thursdays, before and after each watering events. Soil samples were weighed, dried at 60°C for 48 h and weighed again to determine their water content. After six weeks of incubation, detritivores were collected, counted and weighed, and remaining leaf litter was collected carefully, dried at 60°C for 48 h, and weighed. All decomposed litter samples and five samples of initial litter were ground with a disc mill and analyzed for C concentration with a flash CHN Elemental Analyser (Flash Smart, ThermoScientific). The percentage of C lost after the incubation was calculated as [(M_i_ × C_i_ – M_f_ × C_f_) / (M_i_ × C_i_)] × 100 where M_i_ and M_f_ are the initial and final 60°C dry masses, respectively, and C_i_ and C_f_ are the initial and final C concentrations, respectively. We used litter C loss rather than total litter mass loss to correct for inorganic contamination of leaf litter retrieved from mesocosms where they were in direct contact with soil. To isolate detritivore from microorganism effects on total C loss, we subtracted the C loss without detritivore from the C loss with detritivores. This detritivore-driven litter C loss was then corrected for slight differences in detritivore mass among mesocosms by dividing it by the detritivore mass for each mesocosm (average throughout the incubation) and multiplying it by the mean detritivore mass across all mesocosms with detritivores (0.596 g). For mesocosms including both millipedes and isopods, the relative mixing effect was calculated as [(observed value-expected value)/expected value] × 100 for each rainfall treatment separately, where the expected value is the mean of litter C loss driven by millipede and isopods separately, in monospecific treatments. Fourteen mesocosms for which we noted anomalies upon harvest (e.g., presence of snails or lower numbers of detritivores than initially added) were not used for response variable calculations and further data analyses. These removed mesocosms were relatively equally distributed among rainfall treatments (5 for the low quantity and frequency treatment; 3 for all other treatments).

We used two-way ANOVAs to evaluate the effect of rainfall quantity and frequency and their interaction on (1) soil moisture, on the contribution of (2) millipedes, (3) isopods, (4) millipedes and isopods to litter C loss, and on (5) the relative effect of mixing millipedes and isopods. To determine differences in detritivore-driven decomposition among detritivore treatments across all rainfall treatments, we evaluated the effect of detritivore community composition on detritivore-driven decomposition with a one-way ANOVA. For all tests, Tukey HSD post hoc mean comparisons identified difference among treatments. Student’s *t* tests were used to determine if the relative mixing effect for each rainfall treatment was significantly different from zero. All data were checked for normal distribution and homoscedasticity of residuals. All statistical analyses were performed using R version 3.6.1.

## Results

The exclusion of natural rainfall and application of simulated rainfall according to four treatments (current rainfall vs. 50% reduction, each delivered at current frequency vs. 50% reduction) led to variable mean soil moistures throughout the six-week incubation, ranging from 0.25 g H_2_O g soil^-1^ for the low rainfall quantity treatment delivered at low frequency, to 0.31 g H_2_O g soil^-1^ for the high rainfall quantity treatment delivered at high frequency (Fig. 1). Halving rainfall quantity and frequency led to significant relative decreases in mean soil moisture by 7.8% (P < 0.05) and 13.1% (P < 0.001; Fig. 1).

**Figure 1:**
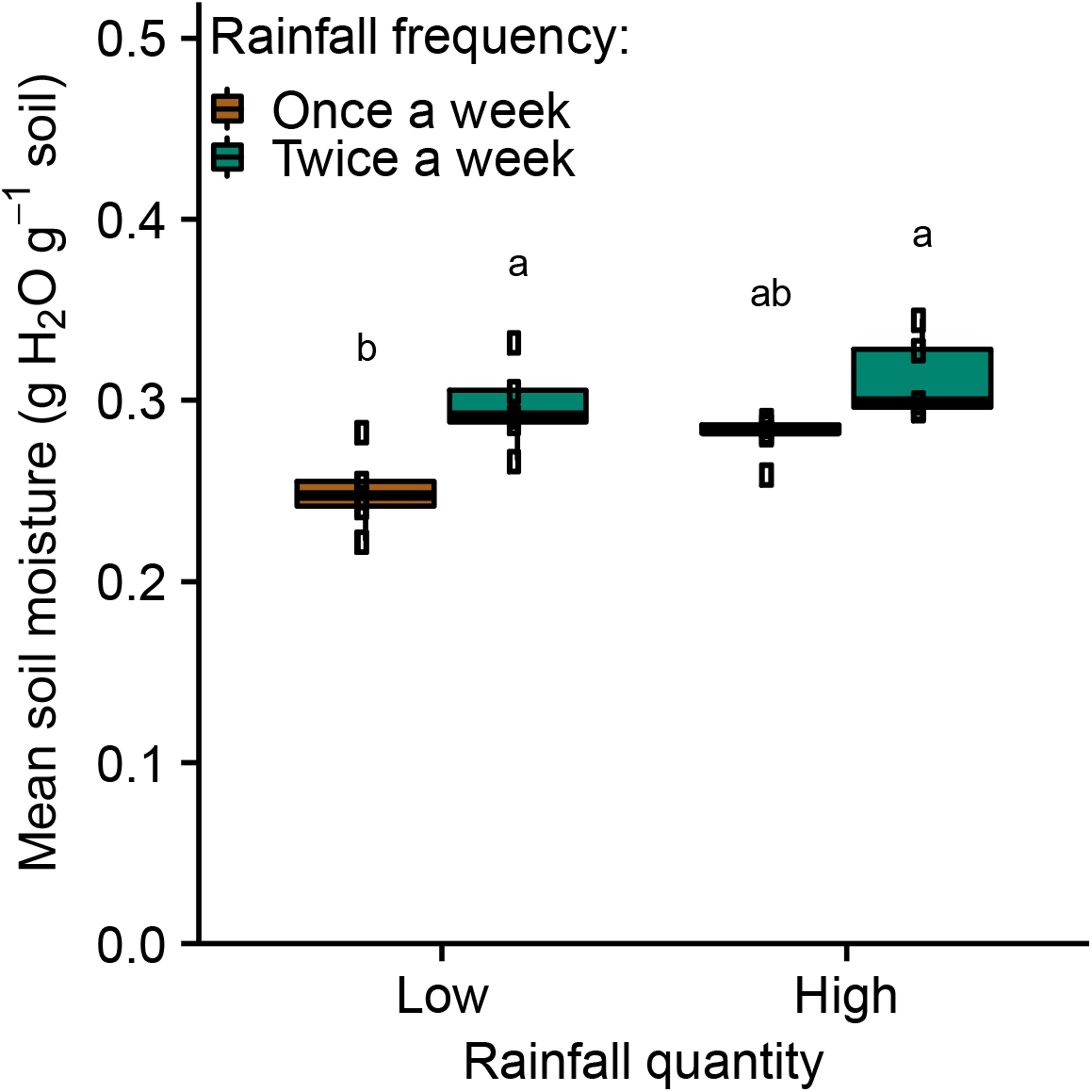
Mean moisture of the topsoil (top 1 cm) throughout the six-week incubation, under two different rainfall quantities (18 and 36 mm/month) and two different rainfall frequencies (one or two pulses per week). Lowercase letters indicate significant differences among rainfall treatments (Tukey HSD tests).

After six weeks of incubation in the field, millipedes contributed significantly more to decomposition than isopods (P < 0.001), with an average detritivore-driven litter C loss of 26.2% for millipedes compared to 10.0% for isopods (Fig. 2), across rainfall treatments. In monospecific treatments, detritivore-driven litter C loss was not significantly affected by rainfall frequency (P = 0.79 and P = 0.76 for millipede- and isopod-driven litter C loss, respectively). They were also not affected by rainfall quantity, although we observed a marginally significant reduction in millipede-driven litter C loss under low rainfall quantity treatments (P = 0.08 and P = 0.62 for millipede- and isopod-driven litter C loss, respectively). When millipedes and isopods cooccurred, their combined contribution to litter C loss reached 22.1%, i.e., a level similar to the contribution of millipedes to litter C loss, but significantly higher than that of isopods (P < 0.001). In contrast to monospecific treatments, detritivore-driven litter C loss was negatively affected by halving rainfall quantity (P < 0.05) but not frequency (P = 0.57) and ranged from 15.3% for the low rainfall quantity treatment delivered at low frequency to 27.4% for the high rainfall quantity treatment delivered at low frequency.

**Figure 2:**
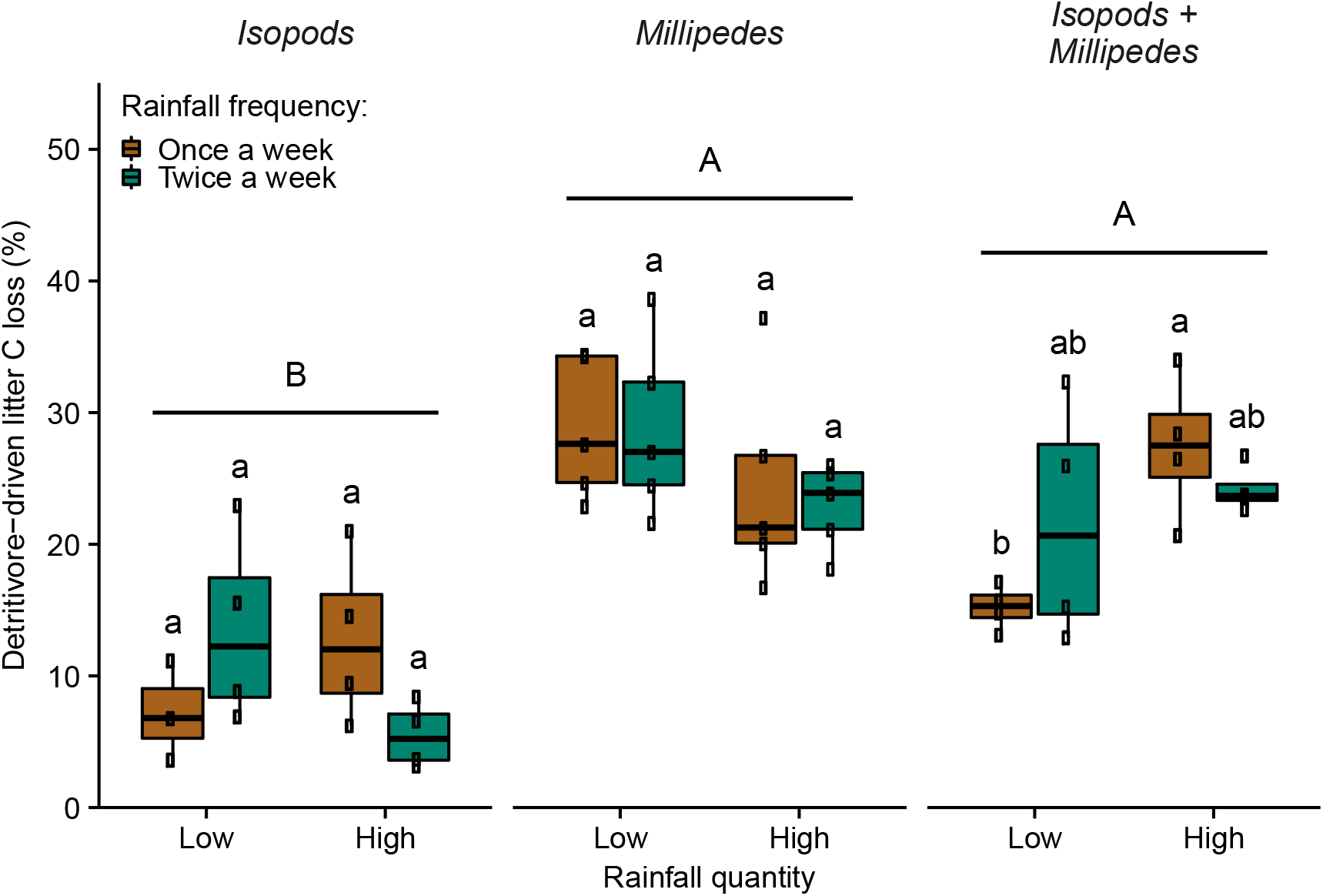
Percentage of litter carbon (C) loss driven by isopods, millipedes, as well as isopods and millipedes, after a six-week incubation under two different rainfall quantities (18 and 36 mm/month) and two different rainfall frequencies (one or two pulses per week). Uppercase letters indicate significant differences among detritivore treatments, while lowercase letters indicate, for each detritivore treatment, significant differences among rainfall treatments (Tukey HSD tests).

The relative effect of mixing isopods and millipedes, i.e., the relative difference between observed values of detritivore-driven litter C loss when millipedes and isopods cooccurred and predicted values based on the respective monospecific detritivore treatments, ranged from −15.4% for the low rainfall quantity treatment delivered at low frequency to +70.1% for the high rainfall quantity treatment delivered at high frequency (Fig. 3). The mixing effect was negatively affected by halving rainfall quantity (P < 0.001) but not significantly affected by halving rainfall frequency (P = 0.16). Except for the low quantity and high frequency treatment, relative mixing effects significantly differed from zero (Fig. 3).

**Figure 3:**
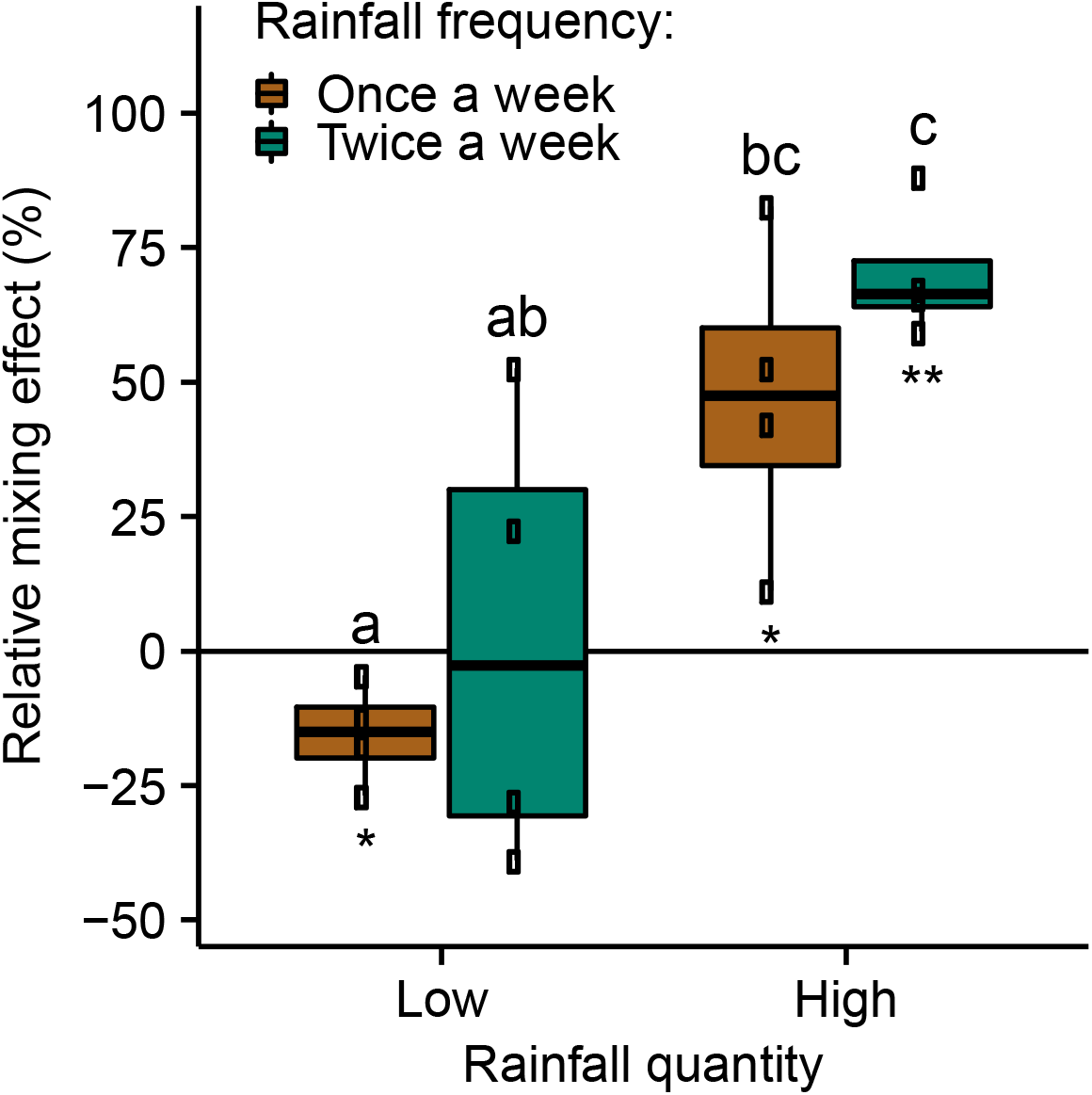
Relative difference between the observed detritivore-driven litter C loss for the millipede-isopod mixture and the expected value based on monospecific treatments after a six-week incubation under two different rainfall quantities (18 and 36 mm/month) and two different rainfall frequencies (one or two pulses per week). Letters indicate differences among rainfall treatments (Tukey HSD tests). Asterisks indicate a relative mixing effect significantly different from 0 (^*^: P < 0.05; ^**^: P < 0.01).

## Discussion

The lack of response of either millipede- or isopod-driven decomposition to halving rainfall quantity supports the view that detritivore feeding activities are largely insensitive to changes in rainfall quantity. This concords with previous studies that reported no response of detritivore-driven decomposition for the isopod species studied therein (Joly et al. 2019) and for another millipede species (Coulis et al. 2013) to altered rainfall patterns and suggests the existence of morphological and/or behavioral desiccation resistance in these detritivore species. Our results indicate that the pattern observed in these studies performed under controlled laboratory conditions, also applies under field conditions. Furthermore, while these previous studies used detritivores collected from drought-prone ecosystems (dryland and Mediterranean shrubland, respectively), the use of detritivores from temperate ecosystems which are not commonly exposed to drought (Scottish Lowlands) in our study suggests that this lack of sensitivity to altered rainfall patterns reflects species rather than population adaptations. It is important to note, though, that despite halving rainfall quantity, we only reported a limited yet significant reduction of topsoil moisture (−7.8%), which may explain the lack of detritivore response to reduced rainfall. Such limited changes in soil moisture suggests that mechanisms other than rainfall quantity/frequency control soil moisture, with water run-off, infiltration and evaporation potentially limiting topsoil moisture under high rainfall quantity, and with air humidity, dew and capillarity potentially limiting topsoil desiccation under low rainfall quantity. On the other hand, halving rainfall frequency induced a larger reduction in topsoil moisture (−13.1%) but also had no effect on either millipede- or isopod-driven decomposition, unlike the expectation in our first hypothesis. This absence of response to even lower topsoil moisture echoes the general insensitivity of these detritivores to drought, but contrasts with the previous findings that isopod-driven decomposition increases with decreasing rainfall frequency (Joly et al. 2019). This finding from a dryland study reflected higher detritivore activities with increasing alternation of dry and moist conditions, suggesting a compensatory feeding in moist periods to satisfy water requirement. In our present study in mesic conditions, the range of litter and soil moisture may not have been sufficient to induce such a response. In turn, strong differences in detritivore-driven decomposition between millipede and isopod treatments suggest that detritivore community composition rather than rainfall patterns controls detritivore-driven decomposition. Such differences may be due to distinct consumption rates between detritivore species, and the higher detritivore-driven decomposition for the millipede compared to the isopod species is in line with previously reported differences in consumption rate for these two species (Coulis et al. 2015). While altered rainfall patterns may have limited effects on detritivore-driven decomposition, effects on detritivore community composition (Zimmer 2004) may have important consequences for decomposition. Furthermore, there is a strong need to consider multiple global change factors such as combined changes in temperature and rainfall that may have interactive effects on the detritivore-driven decomposition (Thakur et al. 2018).

A key result of our study is that interactions between detritivore species markedly varied depending on rainfall pattern with the effect of mixing millipedes and isopods on decomposition ranging from antagonistic (−15.4%) at low levels of rainfall quantity and frequency to synergistic (+70.1%) at high levels of rainfall quantity and frequency (Fig. 3), contrary to our second hypothesis. Such detritivore diversity effects were previously reported (Heemsbergen et al. 2004, DeOliveira et al. 2010), but this aspect of diversity on decomposition remains poorly investigated compared with litter diversity (Gessner et al. 2010, Handa et al. 2014). Although the contribution of isopods and millipedes together to decomposition did not exceed that of millipedes only, our results still suggest that important interactions between detritivores may be at play. Antagonistic effects could be explained by interspecific competition while synergistic effects could result from resource partitioning (use of different litter parts by different detritivore species) or facilitation if litter consumption by one species facilitates litter consumption for the other (Gessner et al. 2010). Importantly, our study suggests that the nature and magnitude of such interactions are strongly dependent on environmental conditions, as evidenced by the change from large positive interactions to non-additive or even negative interactions associated with the reduction in rainfall quantity and frequency. This could be driven by soil or litter moisture (not monitored in this study), with negative interactions decreasing and positive interactions increasing with greater moisture availability. This finding from our field experiment contrasts with laboratory studies that reported no response of detritivore interactions to changes in moisture treatments (Collison et al. 2013, Coulis et al. 2015). Such discrepancies may be due to the consideration of a reduction in rainfall frequency only in Coulis et al. (2015), or due to a lower study duration (two weeks) in Collison et al. (2013). More generally, our results do not support the stress-gradient hypothesis (Bertness and Callaway 1994, He et al. 2013) and suggest that rather than mitigating predicted changes in climate change (“insurance hypothesis”, Yachi and Loreau 1999) positive interspecific interactions on ecosystem functioning may be vulnerable to predicted changes in climate change. Understanding the mechanisms underpinning these detritivore interactions and their sensitivity to rainfall, at larger temporal, spatial and phylogenetic scales thus represents an urgent research avenue.

Collectively, our results indicate that the feeding activities of the detritivore species *Armadillidium vulgare* and *Glomeris marginata* are relatively insensitive to changes in rainfall patterns, but that their interspecific interactions may switch from synergistic to antagonist under reduced rainfall quantity and frequency. Such a reduction in the contribution of diverse detritivore communities to decomposition under future rainfall patterns could slow carbon release from litter. This could have important consequences for soil carbon storage, but an improved understanding of detritivore contribution to soil organic matter formation has yet to emerge (Joly et al. 2020) to predict the consequences on global carbon cycling. However, our focus on a single and rather recalcitrant litter prevents generalizing our findings to litter of different quality. Furthermore, although these species are common across Europe and cooccur in a wide range of ecosystems, our findings may not extend to other detritivore species and their interactions. Evaluating the response of diverse detritivore species with different phylogenetic origins and distribution, and their interactions will be necessary to identify general patterns in the climatic sensitivity of detritivores, thereby better predicting the consequences of climate change on decomposition. To this end, research efforts aiming to identify morphological traits underpinning such sensitivities appear particularly promising (e.g. Dias et al. (2013) for isopods).

## Acknowledgments

We thank Jean-François David and one reviewer for helpful comments and Ian Washbourne for laboratory assistance. This work was funded by a British Ecological Society grant (SR18/1215) and by a UK Natural Environment Research Council grant (NE/P011098/1).

